# Physical Effort Pre-Crastination Determines Preference in an Isometric Task

**DOI:** 10.1101/2024.01.20.576156

**Authors:** Chadwick M. Healy, Alaa A. Ahmed

**Affiliations:** Biomedical Engineering Program, University of Colorado, Boulder, CO, USA; Integrative Physiology Department, University of Colorado, Boulder, CO, USA; Mechanical Engineering Department, University of Colorado, Boulder, CO, USA

**Keywords:** energetic cost, isometric force, pre-crastination, temporal discounting, vigor

## Abstract

How the brain decides when to invest effort is a central question in neuroscience. When asked to walk a mile to a destination, would you choose a path with a hill at the beginning or the end? The traditional view of effort suggests we should be indifferent—all joules are equal so long as it does not interfere with accomplishing the goal. Yet when total joules are equal, the brain’s sensitivity to the temporal profile of effort investment throughout a movement remains poorly understood. Here, we sought to parse out the interaction of time and physical effort by comparing subjective preferences in an isometric arm-pushing task that varied the duration and timing of high and low effort. Subjects were presented with a series of two-alternative forced choices, where they chose the force profile they would rather complete. Subjects preferred to pre-crastinate physical effort but were idiosyncratic about preference for task timing. A model of subjective utility that includes physical effort costs, task costs, and independent temporal sensitivity factors described subject preferences best. Interestingly, deliberation time and response vigor are best described by the same subjective utility model that won for preference, further validating this model of subjective utility. These results suggest physical effort costs are temporally sensitive, with earlier investment of effort preferred to later investment. These findings demonstrate that the representation of effort is based not only on the total energy required but its timing as well, and offer an alternative hypothesis for why animals pre-crastinate in physical tasks.

**NEW & NOTEWORTHY:** This research utilizes a novel paradigm that differentiates between physical effort costs, task costs, and time, where subjects choose between isometric arm-pushing tasks. Here, subjects prefer high physical effort earlier, independent of task timing. We find that deliberation time and response vigor reflect subjective preferences. This research proposes a generalizable subjective utility model that includes independent time-sensitivity factors on physical effort and task costs and offers an alternative hypothesis for why animals may pre-crastinate.

## INTRODUCTION

When asked to walk a mile to a destination, would you choose a path with a hill at the beginning or a path with a hill at the end? Would you ever choose a route with a bigger hill so that you could walk up that hill at a specific time? Traditional models of effort-based decision-making suggest we minimize the total amount of physical effort (Alexander 1997; Hull 1943; Ralston 1958), but these models are indifferent to when periods of high physical effort occur so long as it completes the task. Yet, when posed this simple question, most people have a preference as to when to climb a hill: something the traditional model would not predict. Only recently have investigations shown that humans and animals have a preference for when physical effort costs or task-specific cognitive costs occur (see Rosenbaum and Sauerberger 2022 for a thorough review). Here, we infer the underlying relationship between physical effort, task cost, and time through subjects’ preferences in an isometric arm-pushing task.

Understanding how and when we decide to move (i.e., invest physical effort), is a fundamental question in movement neuroscience. Characterizing how the healthy brain represents movement costs such as physical effort and time is a necessary step in ultimately understanding the changes that occur with age (Lamb et al. 2016) and clinical populations such as Parkinson’s Disease (Mazzoni et al. 2007) or Multiple Sclerosis (Goldman et al. 2008) patients. To date, many questions remain about how the brain subjectively values effort and how effort interacts with other movement costs to determine movement decisions.

In contrast to movement costs, the interaction of reward and time is better defined. With reward-based decision making, reward is temporally discounted, meaning its value decays with time (Haith et al. 2012; Myerson and Green 1995; Shadmehr et al. 2010). Earlier rewards are preferred to later rewards of the same magnitude. Additionally, a reward that requires less effort to obtain is preferred to the same reward which requires more effort (Hartmann et al. 2013; Klein-Flügge et al. 2015). Logically, if reward is discounted by time, and effort is negative reward, then effort costs should be minimized and performed as late as possible. In other words, effort should be procrastinated. Some investigations support this hypothesis, employing a model of utility that includes effort costs as negative reward, with the net reward discounted by time (Bautista et al. 2001; Rigoux and Guigon 2012; Shadmehr et al. 2016). In sum, current models of decision-making optimization would predict effort procrastination: later effort is preferred to earlier effort.

However, other recent observations contradict this: humans and animals prefer to incur some costs earlier. In a study by Rosenbaum et al., participants preferred to complete earlier subgoals at the expense of extra physical effort (Rosenbaum et al. 2014), coining the term *pre-crastination* to describe the subjects’ alacritous approach towards task completion. In these experiments, subjects were asked to walk down an alley, pick up one of two buckets, and carry it to the end. The positions of the buckets varied relative to one another. Surprisingly, most subjects chose to pick up the bucket closer to them, needing to carry the bucket further and expend more energy than the alternative. Since this initial study, numerous experiments have reaffirmed that subjects tend to pre-crastinate certain tasks (Blinch and DeWinne 2019; Fournier et al. 2019a, 2019b; McBride et al. 2023; Patterson and Kahan 2020; Raghunath et al. 2021; Rosenbaum and Dettling 2023; Schwob et al. 2022; VonderHaar et al. 2019; Wasserman and Brzykcy 2015). Many studies that observe pre-crastination are task-centric, if not purely cognitive, and have credited freeing up cognitive resources as the underlying factor for pre-crastination (VonderHaar et al. 2019).

However, it is possible that it is the physical effort cost itself that is pre-crastinated. Indeed, other task-free costs exist, such as pain, in which subjects pre-crastinate. Research has shown that memory of pain follows the peak-end rule, where, in retrospect, people tend to neglect duration and instead remember the most intense moment of an experience and how that experience ends. In cold water baths (Kahneman et al. 1993), a significant majority of participants preferred the longer, objectively more painful trial that ended at a slightly warmer temperature. Patients’ pain evaluation of a colonoscopy procedure correlated to how it ended (Redelmeier and Kahneman 1996). Additionally, in object manipulation tasks, people adopt awkward, hard-to-control positions at first in order to arrive at comfortable, easy-to-control positions later, essentially pre-crastinating discomfort (Cohen and Rosenbaum 2004). Could physical effort be viewed similarly to the discomfort of pain? If physical effort is valued similarly to discomfort, we might observe subjects pre-crastinating high physical effort so they end on a better note.

In many investigations of pre-crastination, expending higher physical effort is tied to subtask completion. If subjects chose to empty heavier buckets first (Fournier et al. 2019b) or carry more dodgeballs earlier (Rosenbaum and Dettling 2023), is this about pre-crastinating physical effort or about task completion? Here, we design a set of effortful tasks that decouple when high physical effort and subtask completion occur to investigate how these costs might be valued relative to one another.

Collectively, what people prefer regarding the timing of physical effort and task costs is not well characterized. We sought to clarify this interaction by comparing subjective preferences across isometric force generation tasks. We decouple physical effort and task costs in order to tease apart what is truly being pre-crastinated or procrastinated. We test this in a novel motor task yet to be explored in the studies of pre-crastination, which controls for timing, accuracy, and explicit information. Across two sessions, subjects choose between effortful isometric arm-pushing tasks that vary in timing and duration of high effort. From each subject’s preferences, we infer a model of subjective utility used as a basis for these decisions. In a group-wise analysis, we find a generalizable model of subjective utility and clarify the relationship between physical effort, task costs, and the time-sensitivity of these costs.

## MATERIALS AND METHODS

### Participants

Twenty-five right-handed, healthy subjects (mean±s.d., age 24.0±5.5 years, 14 female, 11 male) participated in this study. Subjects were required to be between 18-40 years of age, able to understand English, and right-hand dominant. Exclusion criteria included any major musculoskeletal or ophthalmological impairments, low or sedentary activity level (self-reported), and any neurological or vestibular diseases. All subjects provided informed consent, and the experimental protocol was approved by the Institutional Review Board of the University of Colorado Boulder.

### Experimental Setup and Procedure

Subjects sat in front of a computer monitor (1280×1024 px) and used their right arm to grasp the handle of a robotic manipulandum (Interactive Motion Technologies Shoulder-Elbow Robot, Cambridge, MA, Figure 1A). The handle was secured so it would not move, allowing subjects to isometrically push on the handle with their arm. A force sensor (ATI Industrial Automation, Apex, NC) housed inline with the robot arm was used to measure forces exerted by the subject onto the handle. A monitor mounted in front of the subject displayed interfaces that varied depending on the phase of the experiment. Subjects interacted with the displayed interfaces by exerting a force onto the handle which moved a cursor or changed the length of an arrow on the screen.

**Figure 1.**
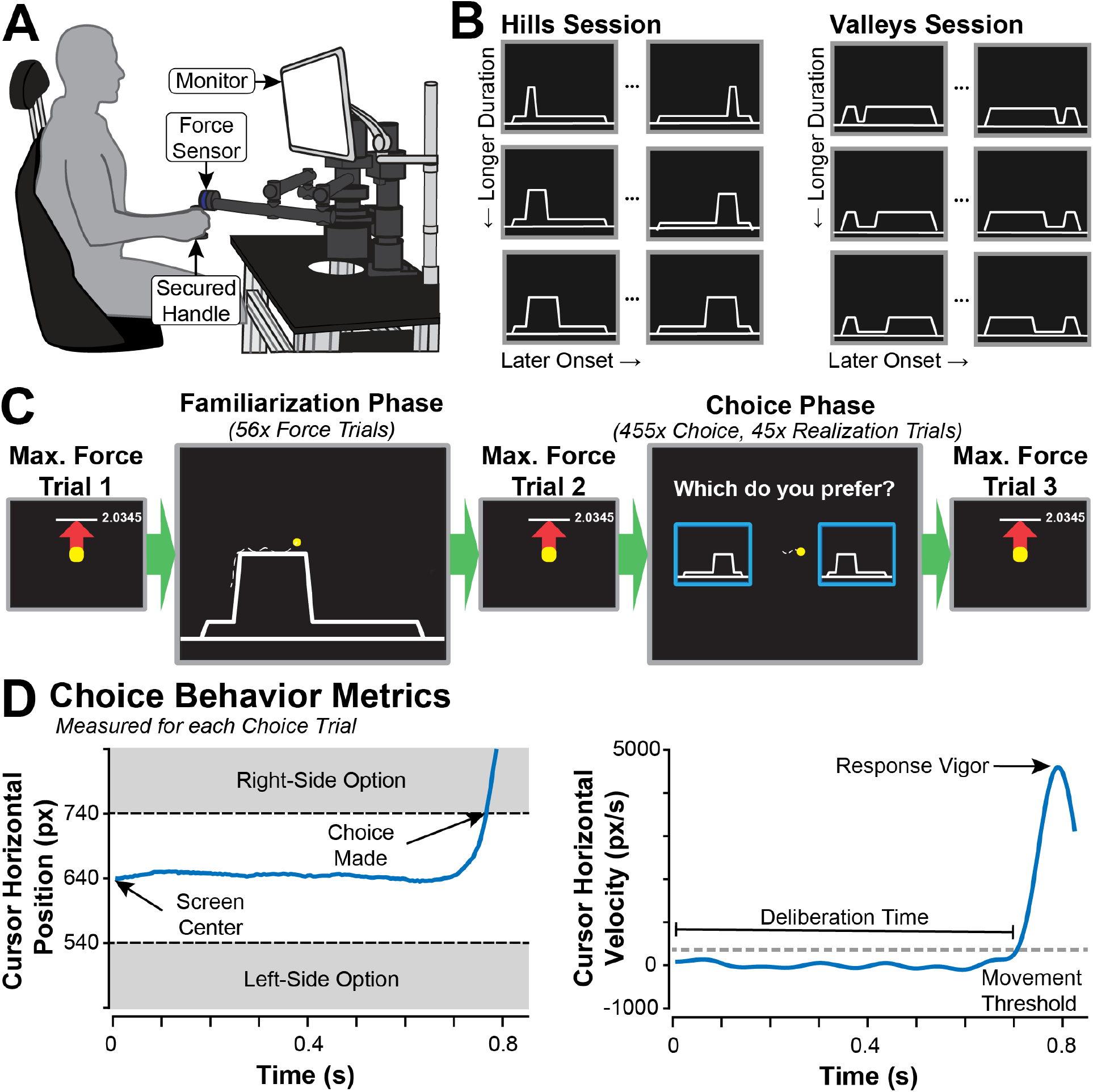
**(A)** Subjects sat in front of a computer monitor and controlled the cursor’s position by exerting force on a handle to a static, secured robotic arm. **(B)** Subjects completed two sessions with identical procedures but different types of force profiles. The Hills Session tested force profiles with brief periods of high effort, while the Valleys Session tested force profiles with brief periods of low effort. Both sessions had 14 unique force profiles determined from combinations of hill/valley duration and onset time. **(C)** Both sessions followed the same five-phase procedure, a Maximum Force Trial was completed three times during a session: at the beginning of the session, between the experiment’s Familiarization Phase and Choice Phase, and at the end of the session. The Familiarization Phase consisted of 56 force trials, and the Choice Phase consisted of 455 choice trials with 45 force realization trials randomly dispersed. **(D)** Data from an example choice trial that shows how deliberation time and response vigor are calculated. (Left) The cursor position starts in the middle of the screen (640 pixels) and then deviates to the right or left, depending on which force profile is preferred. In this example, the subject chooses the right option, moving into the righthand box with a border 100 pixels from the center (740 pixels). (Right) Deliberation time ends when the cursor velocity exceeds a 200 px/sec threshold. Response vigor is calculated as the peak cursor velocity as it moves toward the chosen force profile.

### Experimental Procedure

#### Effort Task Design

Each subject completed two sessions (Hills and Valleys) that followed an identical procedure (Figure 1C). Each session occurred on different days roughly one week apart (mean±s.d.: 8.32±5.91 days; min: 1 day; max: 22 days). The order of sessions was randomized and counterbalanced across subjects.

Each session tested a unique set of force profiles (Figure 1B) which subjects matched using isometric contractions of the shoulder and elbow. The Hills Session tested profiles with brief periods of high effort (i.e., hills) with a low baseline effort required. The Valleys Session tested profiles with a high baseline physical effort and brief periods of low physical effort (i.e., valleys). Hill sessions’ base level of effort was 10% of the subject’s maximum force, with brief hills at 50% of the subject’s maximum force. Valley sessions’ base level of physical effort was 25% of the subject’s maximum force, with brief valleys at 5% of the subject’s maximum force.

For both sessions, we varied the duration of the hill or valley, as well as the time of onset. All force profiles were the same 17 seconds in duration. The hill or valley was one, three, or five seconds in duration and began between 2.5 and 12.5 seconds after the start of the trial. For one-second hills and valleys, we tested six different onset times: 2.5, 4.5, 6.5, 8.5, 10.5 and 12.5 seconds; for three-second hills and valleys, four different onset times: 2.5, 4.5, 8.5, 10.5; and for five-second hills and valleys, four different onset times: 2.5, 4.5, 6.5, 8.5. Not all combinations were possible (e.g., starting a five-second hill at 12.5 seconds exceeds the 17-second duration), resulting in 14 possible combinations of hill/valley onset times and durations per session. A few example profiles for the hill and valley sessions are shown in Figure 1B which visualizes the extremes of duration and onset time.

With this approach, we independently manipulated the timing of two types of costs: *physical effort costs* and *task costs*. Because this is an isometric task, no true mechanical work is performed, so for a given trial, we represent *physical effort costs* as the force-time integral. Hills and valleys were similar in their magnitude of physical effort: trials with three-second hills and three-second valleys had equal force-time integrals. Here, we model *task costs* as a change in state, where these costs are akin to cognitive effort. The attention required to keep force accurate during changes in force level (increasing force to climb a hill or out of a valley, or reducing force to descend from a hill or into a valley) increase cognitive demand and thus task cost. Specific to our paradigm, this is modeled as the time integral of the squared rate of force. So, in terms of Rosenbaum et al.’s bucket task, a hill would actually look like picking up a bucket and setting it back down, where the task costs occur during both picking up and setting down the bucket (Rosenbaum et al. 2014). In our paradigm, valleys incur lower objective task costs than hills, because the rate of change of force is less. Inspecting the force profiles (Figure 1B), a majority of each profile is a flat, constant force level (i.e., no task cost); however, at the beginning and end of a hill or valley, we see a sloped line. With hills, periods of high physical effort and high task cost coincide, making it difficult to tease out which costs the subject may value. However, in valley trials, periods of high task cost now coincide with low physical effort. If we only test one type of session, we cannot separate these costs. For instance, preferring early hills could be a preference for early task costs or early physical effort costs.

However, preferring early hills and late valleys shows a clear preference for early physical effort, because the task is preferred to be performed early in one session and late in the other. With this structure, we can dissociate when the two types of effort occur, and better tease apart if and how temporally sensitive each of these types of costs are.

#### Maximum Force Trial

In each session, subjects were asked to complete three *Maximum Force Trials* throughout the experiment: once at the beginning, once between the *Familiarization Phase* and *Choice Phase,* and once at the conclusion of the experiment. For this task, subjects were asked to make three attempts to push as hard as possible anteriorly against the handle. On the monitor, subjects saw a red arrow grow in height as they pushed harder into the handle, and a white line marked the arrow’s maximum height for the trial (i.e., the maximum force exerted) across all attempts. The maximum force exerted was used to normalize the force experienced across subjects and to assess objective fatigue throughout the experiment. Following the subject’s maximum exertion, they were asked to rate their subjective fatigue on a scale from one to five, where one is “no fatigue” and five is “unable to continue.” Measuring fatigue was essential to ensure that subjects did not fatigue to exhaustion, which could skew decision-making ability.

#### Familiarization Phase

Following the first *Maximum Force Trial*, each session began with the *Familiarization Phase.* Subjects experienced each of the 14 force profiles in a session four times (a total of 56 *Force Trials*). Each force profile type was presented in random order, where it was performed three times in a row. After each force profile type was completed three times, subjects again saw each of the 14 profiles once more in randomized order. Before the final set of 14 trials, subjects were reminded to consider which profile types they preferred. Subjects were given a 10-second rest period every 10 trials. Each trial displayed the force profile as a white line, which subjects were instructed to trace with the cursor. Subjects were given a countdown from three down to zero, displayed numerically and reinforced with audio tones. During the countdown, the yellow cursor remained locked in place at the lefthand side of the force profile. Following the countdown, the yellow cursor started to move horizontally from left to right at a fixed rate. Subjects controlled the cursor’s vertical position by exerting force isometrically against the fixed handle, where a higher anterior force resulted in a higher vertical cursor position. The force trial concluded when the subject’s cursor reached the righthand side of the profile, when the new profile would appear, and the countdown began again. If subjects completed a profile with greater than 7% average error, they were required to repeat it immediately following their prior attempt, however, no subjects exceeded 7% error on any trial. Following the *Familiarization Phase*, subjects completed their second *Maximum Force Trial*.

#### Choice Phase

After the second *Maximum Force Trial*, subjects completed the *Choice Phase*, consisting primarily of *Choice Trials* where subjects performed a two-alternative forced choice task. Unique pairs of the 14 force profiles were presented in a right-left orientation, and subjects were asked to choose which profile they preferred. Each possible combination of profiles for a session (91 unique combinations) was presented five times in random order, totaling 455 choices per session. For a given combination, the order of the profiles (either right or left sides) was randomized. For 10% of choices made (45 per session, randomly dispersed), subjects were required to trace the force profile they selected (*Realization Trials*), reinforcing that their choices had consequences and should be taken seriously. Subjects were given 10-second rest periods every 40 choices.

Each *Choice Trial* began with the subject’s cursor in the middle of the screen with the two options displayed simultaneously with the text prompt “Which do you prefer? Move the cursor to choose.” Deliberation time was unconstrained; subjects were neither forced nor encouraged to choose quickly. Subjects chose which profile type they preferred by exerting a right or left force on the handle moving the cursor to their preferred profile. The horizontal cursor position had the same dynamics as tracing the force profiles. Cursor position away from the middle of the screen was directly proportional to the force they exerted on the handle, normalized to the subject’s maximum force exerted anteriorly. Once the cursor crossed a border of the profile they prefer (±100 px from the center, 12.1% maximum force exerted), an audio cue was given, the profiles disappeared, and a circular target in the center of the screen reappeared with the text “Move back to the center.” After they chose, the cursor was required to be returned to the center of the screen (i.e., no force exerted on the handle) before the next pair of profiles to choose between would appear.

Following a choice trial, 10% of the time, the choice the subject made would need to be performed (*Realization Trial). Realization Trials* were identical to a *Force Trial* in the *Familiarization Phase*, with a countdown, tracing task, and accuracy enforcement. If subjects performed the task with more than 7% average error, they were required to repeat it. Again, no subjects exceeded 7% error on any *Force Realization* trial. Following the *Choice Phase,* participants completed a third and final *Maximum Force Trial*.

#### Data Collection and Processing

A six-axis force sensor embedded in the robot handle measured forces at 200 Hz for all phases of the experiment. Axes of the force sensor were aligned to the robot arm. The angle and orientation of the robot arm were measured and used to rotate forces exerted on the handle to match the orientation of the subject: forces generated in the anterior-posterior direction of the subject moved the cursor vertically, and forces generated to the right or left of the subject moved the cursor right or left on the screen. The handle was secured in a fixed position, so forces were exerted isometrically.

Measured forces were smoothed using a three-point average and used to determine cursor position. Cursor position was recalculated at approximately 10 Hz, and the display refreshed at approximately 60 Hz. Cursor position was linearly mapped to the smoothed forces, scaled to screen size, and normalized by the maximum anterior force exerted for each subject measured during each *Maximum Force Trial,* where 100% of the subject’s maximum force equated to 840 px of deviation from the starting/zero-force point. All data collected were analyzed and processed using custom scripts and built-in functions in MATLAB.

#### Maximum Force Trials

Only the peak force exerted anteriorly from the subject was used during the *Maximum Force Trial*. Maximum force exertions were hand-recorded from the voltage value displayed on the screen. Subjective fatigue ratings were solicited verbally and hand-recorded by the experimenter. These measures were analyzed by experiment phase to investigate if subjects fatigued.

#### Force Trials and Realization Trials

The main measure for *Force Trials* and *Realization Trials* was the accuracy with which the profiles were performed. When subjects traced profiles, forces exerted anteriorly were used to control the cursor’s vertical position, so subjects could visually compare cursor position to the displayed force profile (white lines) in real-time. The difference between the force profile and force exerted was used to calculate the average absolute error, scaled to the proportion of maximum force (82 pixels was approximately 10% of each subject’s maximum force). Error calculations used cursor position data calculated and written at approximately 10 Hz. When the subject had completed tracing the profile, the calculated error was used to determine if the subjects had successfully traced the profile, where any profile completed with more than 7% average error was required to be repeated. The accuracy of each subject’s performance during the *Familiarization Task* was averaged across each unique profile type (four trials each). This metric was used to investigate whether accuracy was a predictor for profile preference, as measured in the *Choice Phase*.

#### Choice Trials

During each *Choice Trial,* we measured the choice made (preferred profile), deliberation time, and choice vigor. These were calculated from the recorded cursor position, forces exerted, and time elapsed for each trial. The cursor position, resulting from forces exerted, was used to determine preference.

Once the cursor crossed the border of the preferred profile (±100 px from the center), the corresponding profile (right or left) was deemed their preferred option. In some infrequent cases, the robot computer would skip to the subsequent trial without letting the subject choose (0.25% of trials). These choices were identified, discarded, and not included in any analyses. Choice behavior measures were determined from the rate of force, which corresponds to vertical cursor velocity (Figure 1D). This was calculated post hoc by regenerating the raw cursor position from measured forces, filtering using a fourth-order Butterworth filter, and then differentiating by time. Deliberation time was calculated as the time of choice appearance until the cursor velocity first exceeded 200 px/sec. Choice vigor was calculated as the maximum cursor speed, which is equivalent to the maximum rate of force normalized to a subject’s maximum force exerted. For the choice behavior analysis, outliers that exceeded the 2.5^th^ and 97.5^th^ percentiles for a given subject were removed. For deliberation time, 3.36% of total choices were not considered in the analysis, and for choice vigor, 2.93% of total choices.

### Statistical Analysis

Our primary methods of analysis were through linear and non-linear models to describe observed behaviors. Below, we detail the structure and measures for each model. Unless specified, results for all models are reported with 95% confidence intervals for parameter estimates.

### A Linear Model of Profile Preference

Our primary variable of interest was the subject’s choice on each trial, as reported by the onscreen cursor’s movement into the preferred option. We first investigated how subjects’ preferences were determined by the variables we intentionally manipulated: duration, onset time, and session type. We coded choices as a binomial dataset*, ChoseLeft*, where 1 indicates the left option was chosen, and 0, the right. We employed a logit-linked binomial generalized linear mixed-effects model to investigate the fixed covariates and interactions of session type (*SessionType*, categorical with two levels: hill or valley), the difference in hill or valley duration (*Δduration*, continuous, seconds), the difference in hill or valley onset time (*Δonset,* continuous, seconds), and *Subject* (categorical) as a random intercept term. The model is summarized in Equation 1 below, where *ChoseLeft_ij_* is the *j*th choice of subject *i*, and *Subject_i_* is normally distributed with mean 0 and variance σ^2^.

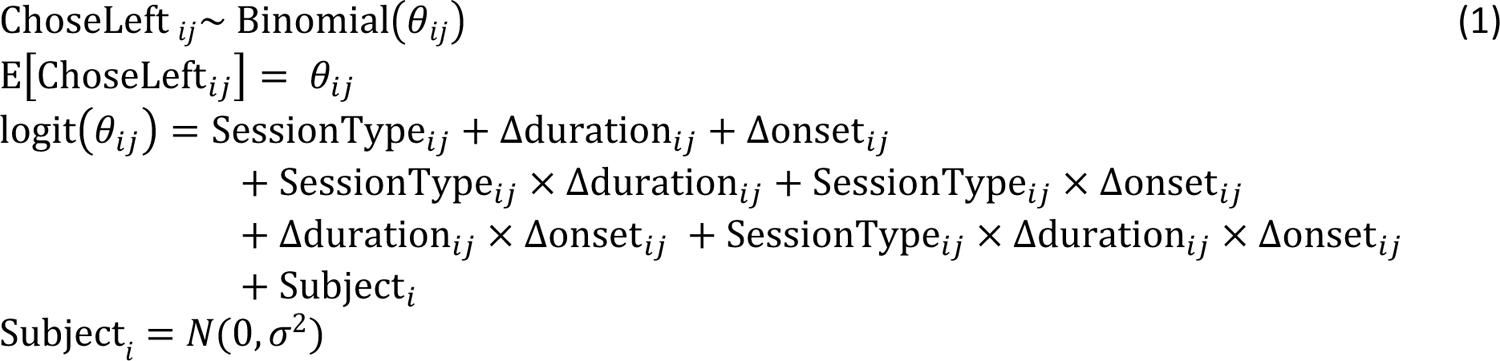

### Profile Preferences Using Models of Subjective Utility

We employed a more complex but more generalizable model for subject preference. Here, choices are modeled as a function of the difference in subjective utility between profile types, where subjective utility is determined from the actual force profile versus time, instead of protocol-specific factors like duration or onset time of a single hill or valley. We compared four different models of subjective utility that incorporate different specific costs (physical effort costs, task costs, and time sensitivity) and their interactions. With this approach, we can infer more specifically what costs are driving subjects’ preferences.

For all models of subjective utility, we quantified the decision-making process as a function of the difference between the subjective utility of each option. The probability of choosing the left profile was structured as a sigmoidal function of the difference between the subjective utility for each choice, where subjective utility was represented by *U* for option *left* or *right,* and a subject-specific temperature parameter,*τ*:

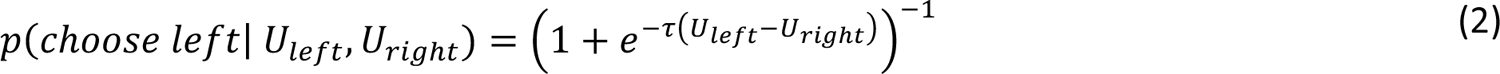

We calculated utility as reward, *R*, minus costs, *E*. In this specific experiment, there was no explicit reward thus no difference in reward across profiles. Mathematically, we modeled reward to be zero resulting in a utility that was simply the negative of the subjective effort, *E*.

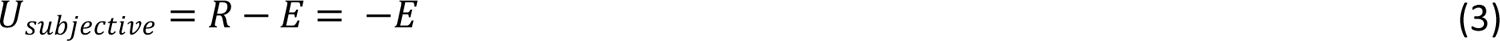

We distinguished between two types of costs: *physical effort* and *task.* Physical effort costs for a given profile were represented as force normalized to a subject’s maximum and the force-time integral of a one-second hill profile, *K_physical_* (Equation 4). Task costs were represented by the rate of force squared normalized to a subject’s maximum, and the force-time integral of a one-second hill profile, *K_task_* (Equation 5). This normalization was important so both types of effort were the same order of magnitude. Examples of physical effort costs and task costs in different profile types are shown in Figure 2.

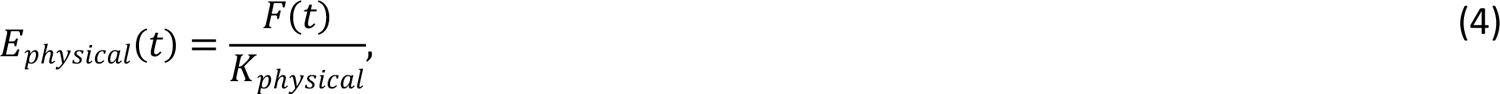

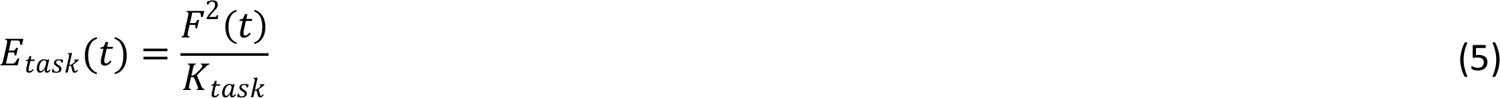

**Figure 2.**
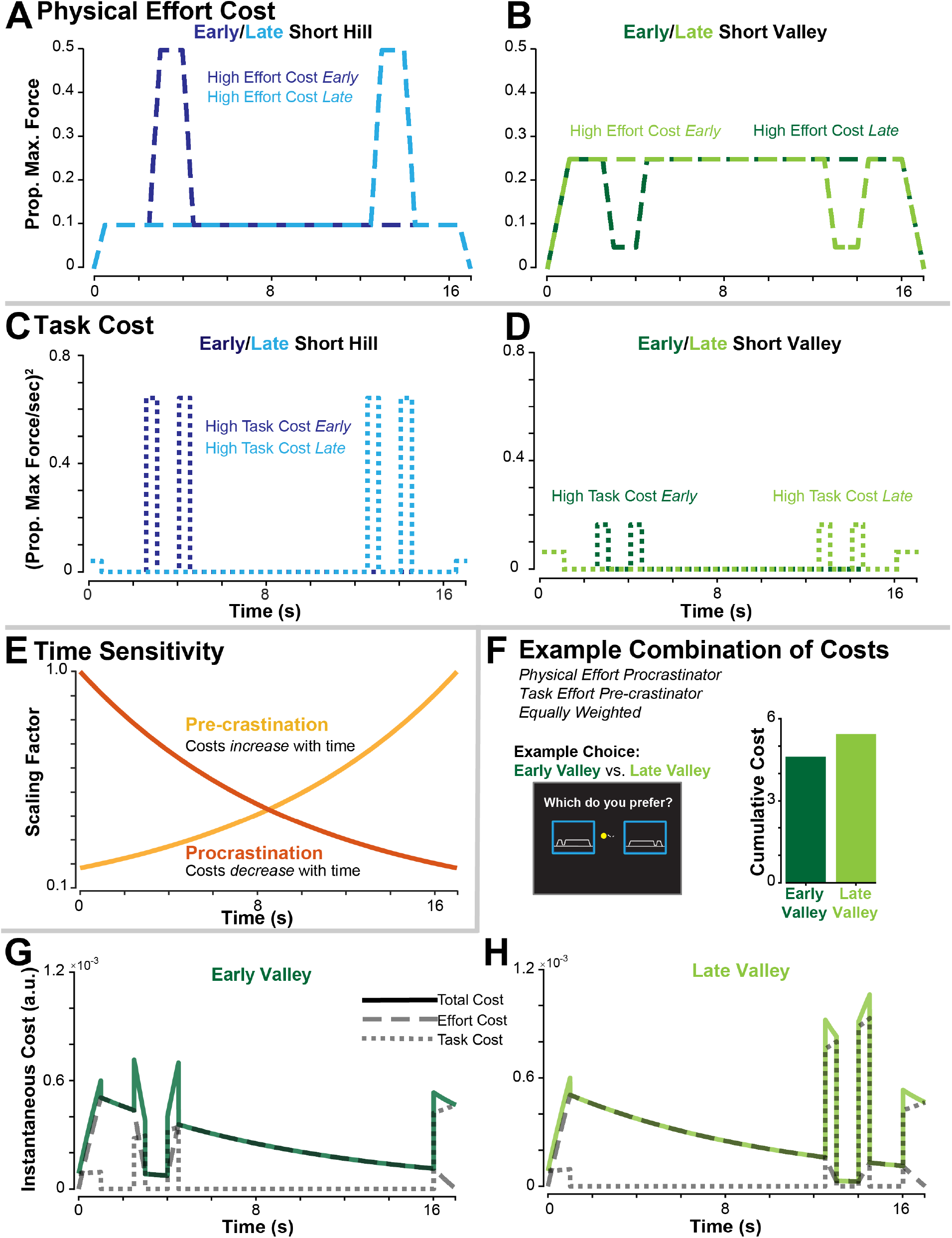
How subjective effort costs are calculated from the force profiles is shown, including an example choice. Physical effort cost (dashed lines) is represented as the force-time integral. **(A)** An early, short hill (dark blue) has high physical effort cost early and low physical effort cost late, and a late, short hill (light blue) has high physical effort cost late and low physical effort cost early. **(B)** An early, short valley (dark green) has low physical effort cost early and high physical effort cost late, and a late, short valley (light green) has high physical effort cost early and low physical effort cost late. Task costs (dotted lines), represented as rate of change of force squared, corresponds to when the hill or valley occur. For both **(C)** early short hills (dark blue) and **(D)** early short valleys (dark green), high task costs occur early, while late short hills (light blue) and late short valleys both have high tasks costs late. **(E)** Time sensitivity is represented with two possible categories: pre-crastination (yellow) and procrastination (orange). This cost function multiplies with physical effort costs or task costs to combine into a time sensitivity for when those costs occur. For pre-crastination, costs increase with time, and for procrastinators, costs decrease with time. Using this structure to represent effort costs, they combine per the various utility functions (Equations 7-10). Using the fourth model of utility (Equation 10), an example of how the costs combine is illustrated. In this case, we show how someone who is an effort cost procrastinator, task cost pre-crastinator, with equal weights on types of costs, might experience an early valley **(G)** versus a late valley **(H)**. Total costs are shown in bold color with the effort costs shown as a dashed line, and task costs, a dotted line. Each profile’s cost is integrated, and whichever profile has the lower cumulative cost is the preferred option. **(F)** In this case, we see the early valley (dark green) has lower cumulative cost and is the preferred option.

We modeled the cost of time as a function that could be multiplied with effort costs to exhibit temporal sensitivity for those costs (Figure 2E). Previous studies model temporal sensitivity as an exponential (Rigoux and Guigon 2012) or hyperbolic discounting function (Haith et al. 2012). For this study, we used a piecewise exponential formulation (Equation 6), where γ is the sensitivity factor, *t* is instantaneous time, and *T* is the end time (in our case, always 17 seconds). A positive γ value is a cost procrastinator, where costs further in the future incur a smaller penalty than costs closer to the present. Alternatively, a negative γ value is a cost pre-crastinator, where costs in the future are a larger consequence than costs expended in the near term. Using the piecewise function structure allows for the effect of positive and negative γ-values was equally distributed from zero.

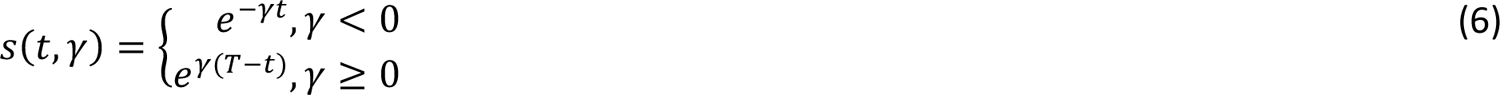

We explore four different subjective utility models, where each is different cost combinations from Equations 4, 5, and 6. First, we first considered a traditional model of effort: not time-sensitive and simply the force-time integral of the profile (Equation 7). This first model of subjective utility, *U_1_*, predicts a preference for shorter hills and longer valleys but indifference as to when those hills and valleys occur.

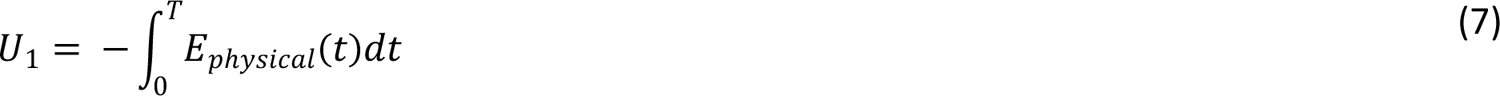

Next, we explored a second model of subjective utility that incorporates time sensitivity and physical effort (Equation 8). This second model, *U_2_*, allowed predictions of subjects who prefer when periods of physical effort occur. For example, this model could describe subjects who preferred early hills and late valleys and classify them as physical effort pre-crastinators.

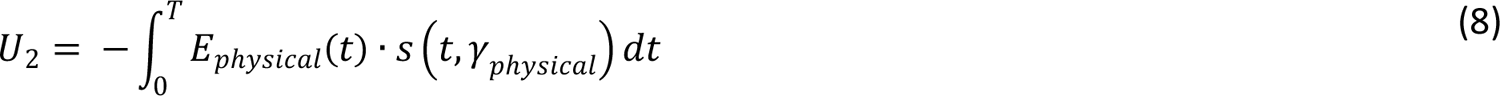

Next, we considered a third model, *U_3_*, that incorporates task costs, physical effort costs, and time sensitivity, γ*_combined_*, (Equation 9). However, we constrained this model so that both are temporally sensitive in the same way. A weighting parameter, α, varies between 0 and 1, which shifts the importance of physical effort versus task costs. This model could describe subjects who prefer early hills and early valleys, classifying them as pre-crastinators of both physical effort and task costs with a stronger weighting towards task costs.

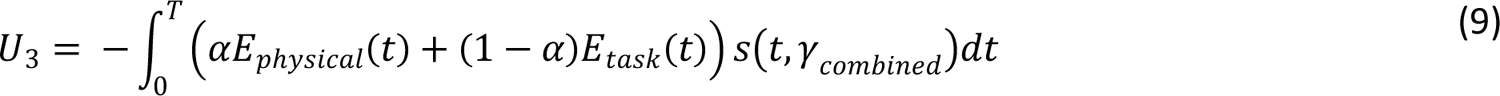

Finally, we considered a model of utility, *U_4_*, that includes physical effort and task costs with independent temporal-sensitivity factors, γ*_physical_* and γ*_task_* (Equation 10). This model offers an alternative explanation to *U_3_* for a subject who prefers early hills and early valleys: they could be task cost pre-crastinators and physical effort procrastinators, and still care more about task costs.

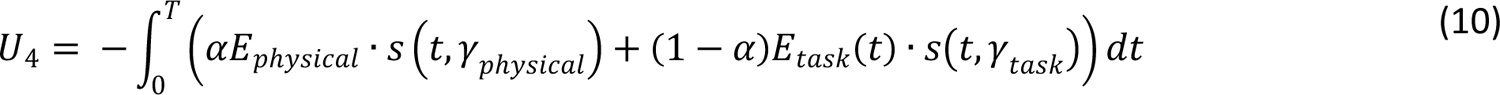

### Predicting Subject Preferences with Utility Functions

The models summarized above are nonlinear, thus we employed MATLAB’s constrained minimization function, *fmincon*, which is designed for nonlinear optimization problems. For each subject, we minimized an objective function (Equation 11), which is the negative log-likelihood of the probabilistic model’s prediction for each subject’s choices (Equation 2). For the total number of valid trials, *N*, this function compares if the subject chooses the left option (binary outcome variable, *y*) to the model’s predicted choice, *p*, for a given profile combination. For a given utility model, each subject was fit independently. Parameters were constrained but allowed to vary within set bounds. The minimizing solution was obtained for each fit by comparing results from multiple restarts with randomized initial parameter values. The solution with the smallest objective function (i.e., maximum likelihood) was chosen. Parameters for each subject and model were then collated and compared, and the aggregate model performance across subjects was analyzed to see if a single model best described subject choice data.

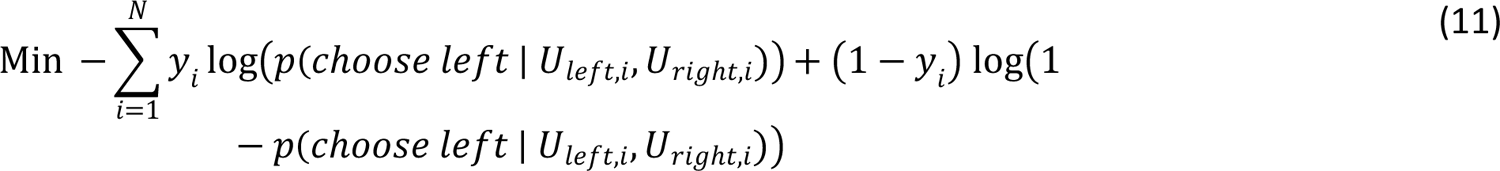

### Utility Model Selection and Estimating Population Parameters

The most likely model was determined using Bayesian model selection (BMS) (Rigoux et al. 2014; Stephan et al. 2009). Using the calculated log-likelihood for each subject-model pair, the Akaike Information Criteria (AIC) was calculated. Model posterior means and protected exceedance probabilities (pxp) were computed using AICs with the Statistical Parametric Mapping software (SPM, version 12). Protected exceedance probability is the probability that a given model is more likely, tested against the null hypothesis that all models are equally likely. The model with the highest pxp was selected as the winning model.

From the winning model, we estimated population distribution for each parameter by bootstrapping each set of fit parameters 10000 times with replacement. From these confidence intervals, we determined how the population may be weighting the various costs of physical effort costs, task costs, and time.

### Choice Behavior and Preference

Both deliberation time and choice vigor were fit using linear mixed-effects models. For each model of utility (Equations 7-10) and each subject, the absolute difference in subjective utility was min-max normalized using Equation 12. The normalized difference in subjective utility was used as the independent variable to explain the dependent variables deliberation time and choice vigor.

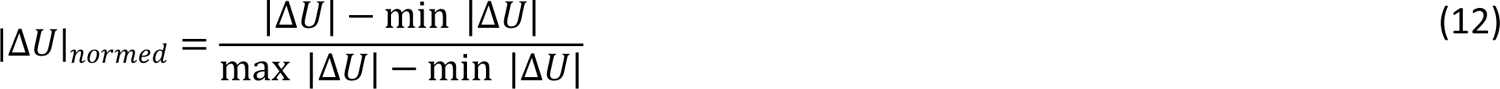

Deliberation time was modeled using a simple linear mixed-effects model with a Gaussian response variable (Equation 13). We investigated the fixed effect of the normalized difference in subjective utility (|*ΔU*|*_normed_*, continuous from 0 to 1), a fixed intercept term, and a random effect of subject (*Subject*, categorical) on the intercept. In the model, *DelibTime_ij_* is the deliberation time of the *j*th choice of subject *i*, and *Subject_i_* is normally distributed with mean 0 and variance σ^2^.

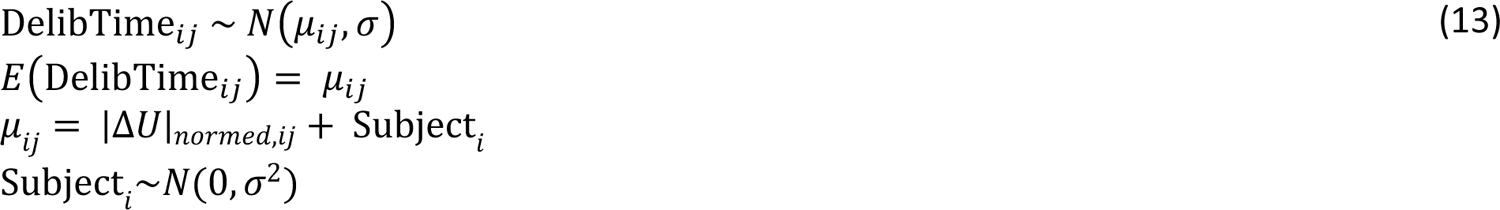

We investigated choice vigor using an identically structured linear mixed-effects model with a Gaussian response variable (Equation 14). Again, we include the fixed effect of the normalized difference in subjective utility (|*ΔU*|*_normed_*, continuous from 0 to 1), a fixed intercept term, and a random effect of subject (*Subject*, categorical) on the intercept. In the model, *ChoiceVigor_ij_* is the vigor of the *j*th choice of subject *i*, and *Subject_i_* is normally distributed with mean 0 and variance σ^2^

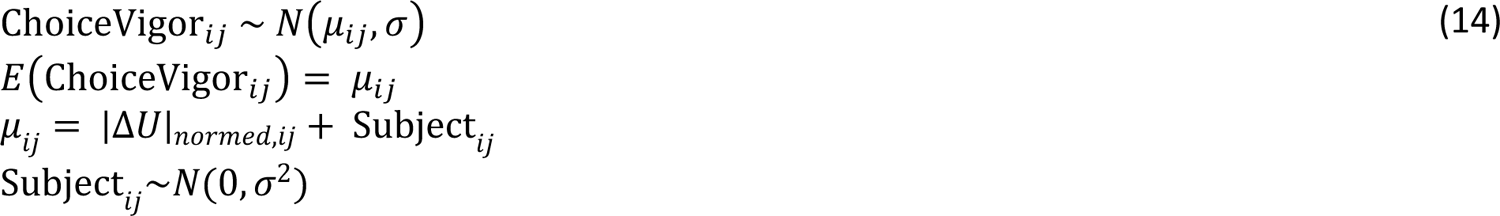

Finally, we tested if deliberation time can be predicted by choice vigor using a linear mixed-effects model (Equation 15). We model deliberation time (*DelibTime*, continuous) as a Gaussian response variable, a fixed effect of choice vigor (*ChoiceVigor,* continuous), a fixed intercept term, and a random effect of subject (*Subject,* continuous) on the intercept.

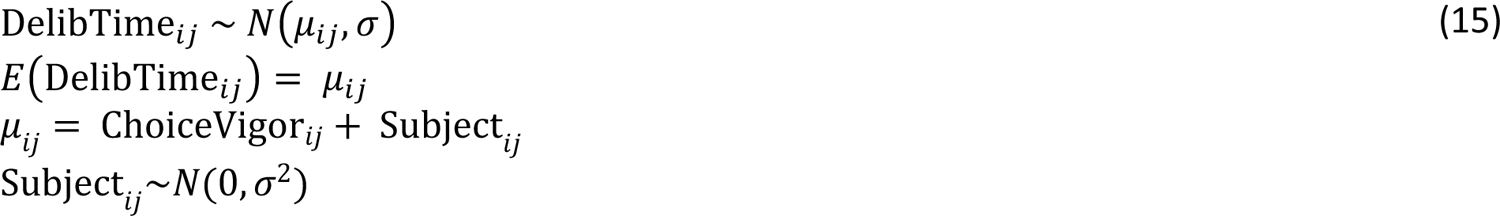

### Accuracy as a Predictor of Preference

We investigated if subject accuracy when performing a profile predicted their preferred profile. Because not all profiles were performed during the *Realization Trials* in the *Choice Phase,* we used the average error for each profile type from the *Force Trials* completed in the *Familiarization Task*. Using the binomial choice data, *ChoseLeft,* we constructed a logit-linked binomial generalized linear mixed-effects model with the fixed effect of difference in error between profile types (*Δerror*, continuous) and a random intercept term for subject (*Subject,* categorical). The model is shown below in Equation 16, where *ChoseLeft_ij_* is the *j*th observation of subject *i*, and *Subject_i_* is normally distributed with mean 0 and variance σ^2^.

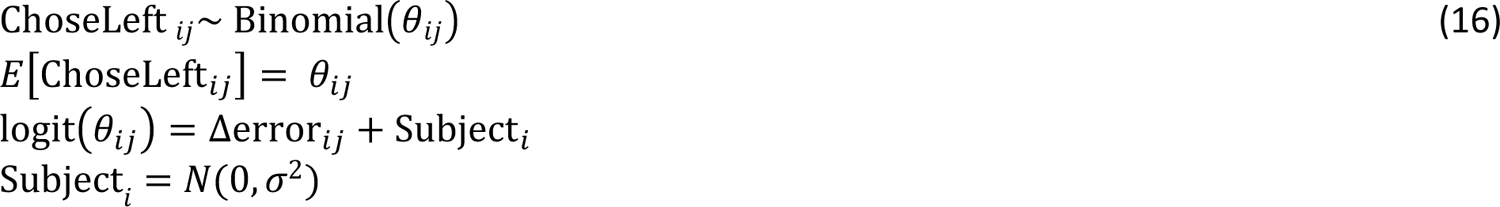

In addition, we investigated the correlation between the difference in error and other predictors investigated in Equation 1 (difference in onset, difference in duration, session type, and their interactions) using a linear mixed-effects model with a random intercept term for subject (Equation 17). This model used the set of all possible choice combinations (91 per session, 182 per subject) rather than the full set of choices used, as with Equations 15 and 16. We investigated the fixed covariates and interactions of session type (*SessionType*, categorical with two levels: hill or valley), the difference in hill or valley duration (*Δduration*, continuous, secs), the difference in hill or valley onset time (*Δonset,* continuous, secs), and *Subject* (categorical) as a random intercept term. The model is summarized in Equation 17 below, where *Δerror_ij_* is the *j*th choice combination of profile types for subject *i*, and *Subject_i_* is normally distributed with mean 0 and variance σ^2^.

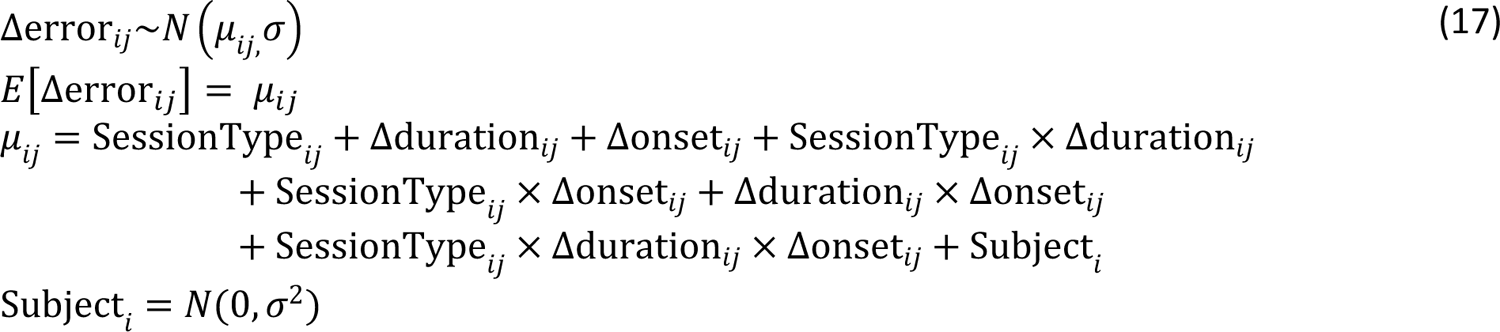

### Subject Fatigue by Experiment Phase

We used the measured maximum force exerted and subjective fatigue rating collected during the *Maximum Force Trial* to determine if subjects fatigued throughout the experiment. We investigated how the two measures of fatigue change across the three measurements (*ExpPhase,* ordinal, three values) and session type (categorical, hill or valley). Each measure used an identical linear mixed-effects model shown in Equation 18, where *Fatigue_ij_* is *j*th observation either the maximum force exerted or the subjective fatigue rating for subject *i*.

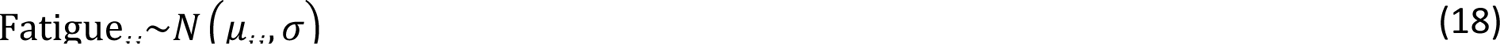

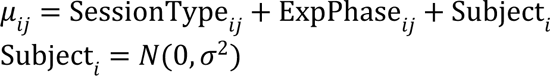

## RESULTS

### Profile Preference

We first investigated underlying factors for subject preferences (Table 1) using a generalized linear mixed-effects model. Notably, we observed a significant effect of duration (mean and 95% confidence interval: β=0.905, [0.872, 0.938], p<0.0001), where the positive coefficient indicates a preference for shorter hills (Figure 3, top left). There was also a significant interaction between duration and session type (β=−1.761, [−1.81, −1.72] p<0.0001), with a reversal of sign demonstrating that preference is shifted towards longer valleys (Figure 3, top right). Interestingly, there is a significant preference for earlier hills (β=0.0448, [0.0338, 0.0558], p<0.0001) and a significant interaction between onset time and session type (β=-0.0922, [−0.107, −0.0768], p<0.0001) that reveals a preference for later valleys. There was also a significant effect of the interaction between duration, onset time, and session type (β=-0.0122, [−0.0217, −2.63×10^-3^], p=0.0124), which suggests there is a stronger preference for later valleys than earlier hills (Figure 3, bottom).

**Figure 3.**
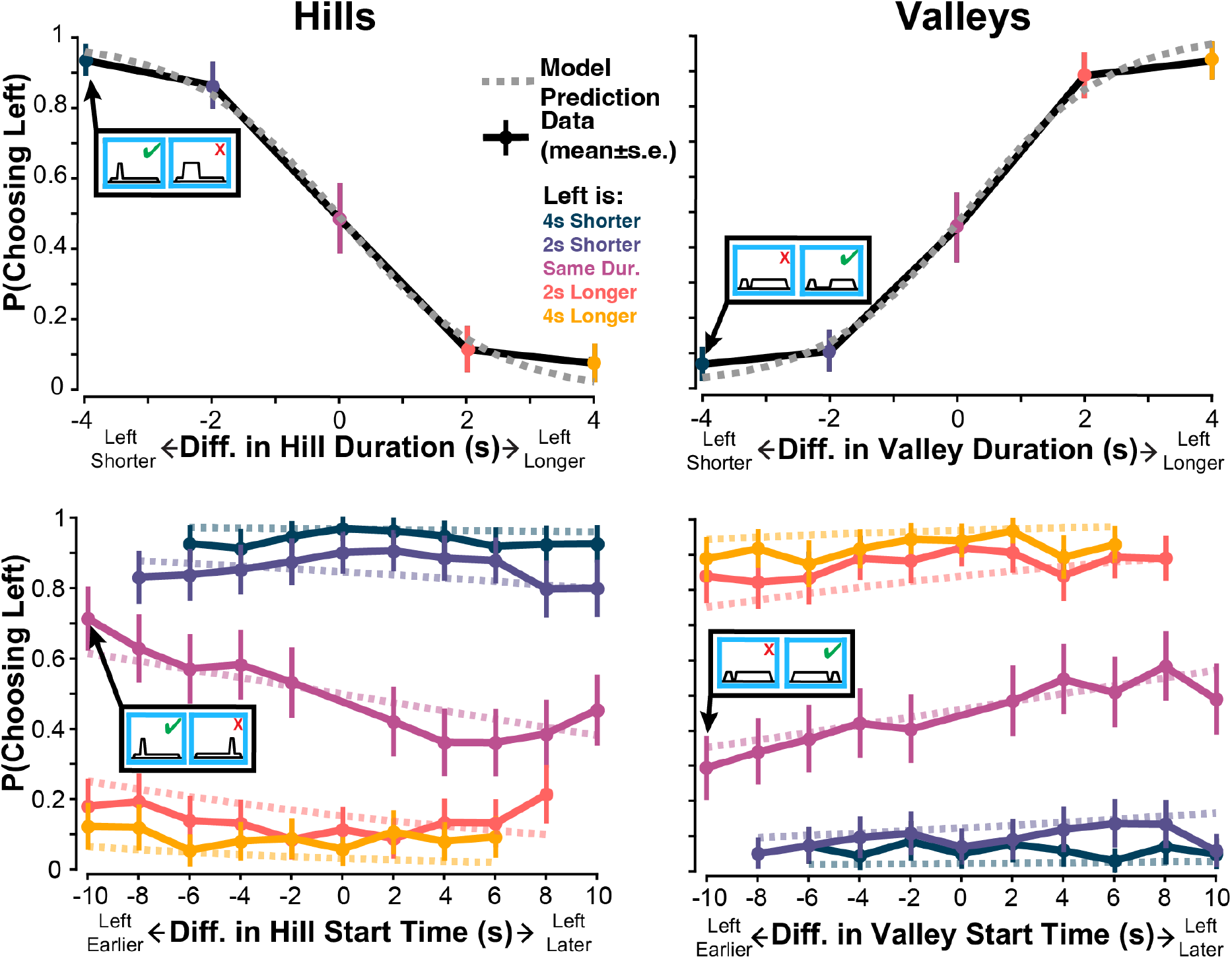
**(Top Row)** The probability of choosing a hill (left) or valley (right), based on the difference in duration between options (left-right). Mean and s.e. choice data are shown with colored dots and bars connected by black lines. The dashed line is the predicted probability from the linear model. Across subjects, there is a strong preference for shorter hills and longer valleys. **(Bottom Row)** The probability of choosing a hill (left) or valley (right) depends on the difference in onset time. Choices are stratified by difference in duration, where blue is four seconds shorter, purple is two seconds shorter, pink is equal duration, orange is two seconds longer, and yellow is four seconds longer. Mean data is shown with dots connected by bold, colored lines, and model predictions are shown as dashed lines. The dominant effect on preference is duration, but when hill or valley duration is equal, there is a higher probability of choosing an earlier hill or a later valley.

**Table 1:** Generalized Linear Mixed-Effects Model of Subject Preference.

| Fixed Effect | Estimate (mean, 95% CI) | P-value |
| --- | --- | --- |
| Intercept | -0.156 [-0.307, -0.00513] | 0.0427 |
| SessionType=Valley | -0.152 [0.0783, 0.225] | <0.0001 |
| Δduration | 0.905 [0.872, 0.938] | <0.0001 |
| Δonset | 0.0448 [0.0338, 0.0558] | <0.0001 |
| Δduration x SessionType=Valley | -1.76 [-1.81, -1.72] | <0.0001 |
| Δonset x SessionType=Valley | -0.0922 [-0.107, -0.0768] | <0.0001 |
| Δduration x Δonset | 4.95x10<sup>-3</sup> [-1.94x10<sup>-3</sup>, 1.18x10<sup>-2</sup>] | 0.15892 |
| Δduration x Δonset x SessionType=Valley | -0.0122 [-0.0217, -2.63x10<sup>-3</sup>] | 0.0124 |

These results make sense: subjects were sensitive to total duration of physical effort and preferred less effort to more effort. Interestingly, they also show that subjects were sensitive to when effort was invested, not just for how long. For effort of similar duration, subjects tended to pre-crastinate and prefer high effort earlier than later (earlier hills and later valleys).

### Preferences Suggest Time Sensitivity of Physical Effort and Task Costs

Subjects’ choices were fit using the various models of subjective utility (Equations 7-10). The four-parameter model of utility (Equation 10, *U_4_*), which includes physical effort costs, task costs, and independent time sensitivity factors, best explained subject choices (Table 2). This model has the highest pxp of 0.986, as determined from the group-level BMS. This model (*U_4_*) best described data from 13 of the 25 subjects compared to 6 subjects for *U_3_*, 3 subjects for *U_2_*, and 2 subjects for *U_1_*.

**Table 2:** Aggregate AIC scores and protected exceedance probabilities (pxp) for choice behavior for each model of subjective utility.

| Model | Number of Parameters | Aggregate AIC | Model Posterior Means | Protected Exceedance Probability |
| --- | --- | --- | --- | --- |
| Force Only ($U_1$) | 1 | 17405 | 0.0429 | $1.79 \times 10^{-3}$ |
| Time Sensitive Force ($U_2$) | 2 | 16444 | 0.139 | $2.73 \times 10^{-3}$ |
| Time Sensitive, Combined Force and Rate of Force ($U_3$) | 3 | 14823 | 0.2141 | $9.39 \times 10^{-3}$ |
| Independent Time Sensitivity, Force and Rate of Force ($U_4$) | 4 | 14389 | 0.6041 | 0.986 |

We investigated the results of the winning model in Figure 4, where we see average choice behavior matches model predicted choice behavior (Figure 4A). Because individual subjects were fit independently, it is also valuable to inspect individual fit performance. Here, we show a particular subject who was classified as a physical effort pre-crastinator and task cost pre-crastinator, with an emphasis on physical effort over task costs (γ*_physical_* = −0.0202; γ*_task_* = −0.524; ⍺=0.999). In Figure 4B, we see the model captures the subject’s choice behavior well as a function of the difference in subjective utility. For this subject, we rank each profile’s subjective utility to intuit what the model predicts the subject cares about. For this particular subject, a physical effort pre-crastinator, we see that early, short hills are most preferable and late, long hills are least preferable. Within valleys, we see that long late valleys are preferable and short early valleys are least preferable. This ranking matches the description of a physical effort pre-crastinator.

**Figure 4:**
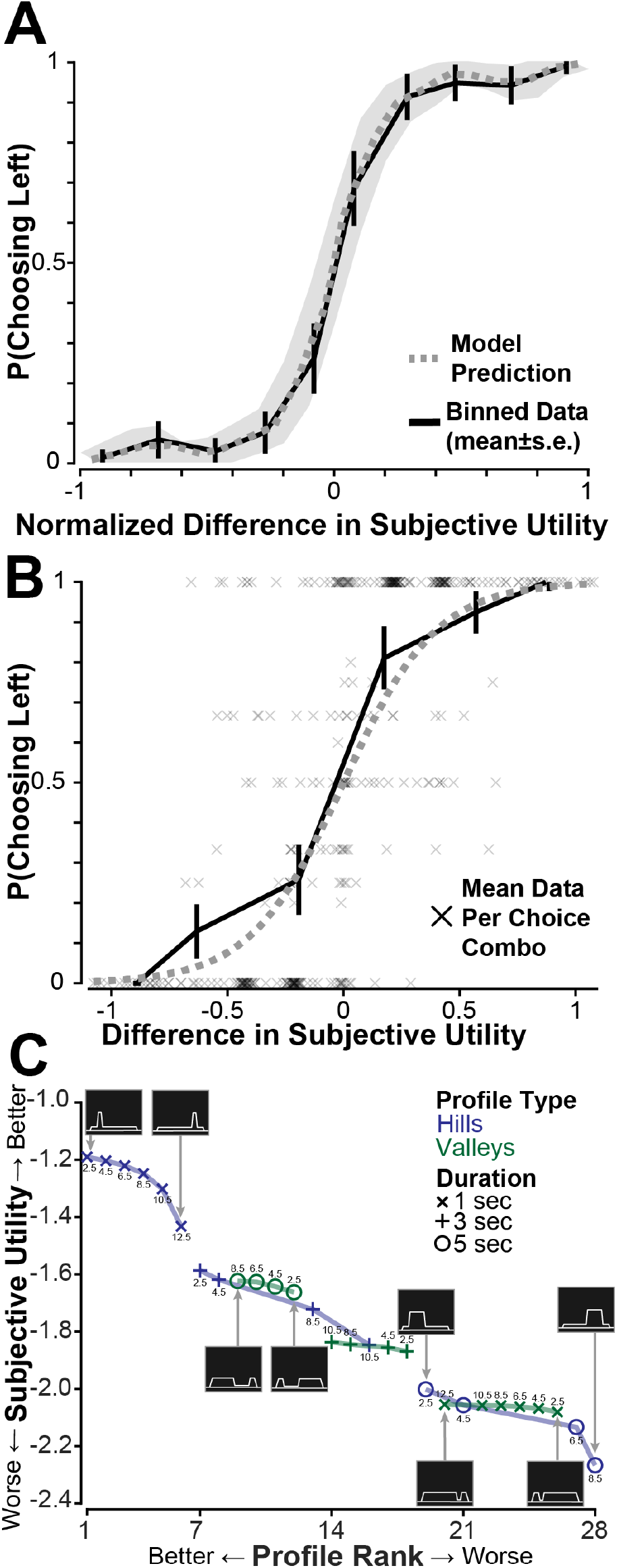
**(A)** Model predicted probability of choosing the left option, given the normalized difference in subjective utility (left-right) for the winning model averaged across subjects. Model predictions with 95% confidence intervals are shown in gray. Choice data was binned and averaged, with mean and standard error bars shown in black. Data and model predictions from a representative subject are shown in **(B)** and **(C).** This subject was characterized as a physical effort pre-crastinator (γ*physical* = −0.0202) and task cost pre-crastinator (γ*task* = −0.524), with a strong emphasis on physical effort over task cost (⍺=0.999) **(B)** Model predicted probability of choosing the left option for the example subject is shown in gray, using subjective utility from the winning model. Mean probability of choosing left for a given combination (two unique profile types and which profile is on the left) is shown as a black x. Binned choice data is averaged, with the mean and standard deviation bars shown in black. **(C)** Using the subjective utility, each profile is ranked against one another. Hill profiles are shown in blue, and Valley profiles are shown in green. The different durations of hills and valleys are shown using different markers per the legend, with the onset time indicated numerically. For this subject, the most preferable option is short, early hills, and the least preferable options are short, early valleys and late, long hills. For this subject, the pattern is consistent, where, for a given duration of hill or valley, preference decreases as high physical effort occurs later.

### Subjects Pre-Crastinate Physical Effort Costs

Investigating parameter estimates for the winning utility model, *U_4_*, we bootstrap each set of parameter values with replacement 10,000 times to estimate the population distribution (Table 3). For time sensitivity factors, a negative value indicates *pre-crastination,* whereas a positive value indicates *procrastination.* For the weighting term, a value of ⍺=0.5 equally values *physical effort* and *task costs*, ⍺=1 values only *physical effort costs*, and ⍺=0 values only *task costs*.

**Table 3:** Bootstrapped parameter values for the winning model of subjective utility.

| Parameter | Confidence Interval (95%) |
| --- | --- |
| Temperature ($\tau$) | $[1.37 \times 10^{-3}, 5.76 \times 10^{-3}]$ |
| Time Sensitivity of Physical Effort Costs ($\gamma_{physical}$) | $[-0.0903, -0.00499]$ |
| Time Sensitivity of Task Costs ($\gamma_{task}$) | $[-0.134, 0.125]$ |
| Weighting Between Physical Effort Costs and Task Costs ($\alpha$) | $[0.234, 0.582]$ |

These results suggest that subjects pre-crastinate physical effort costs (CI: [−0.0903, −0.00499]), where high physical effort is preferred earlier rather than later. Time-sensitivity of task costs is nearly centered at zero (CI: [−0.134, 0.125]), showing no consistent trend across subjects. Weighting between physical effort and task costs (CI: [0.234, 0.582]) shows that subjects tend to value when the task costs occur equally, if not slightly more than physical effort alone. These factors, in combination with *U_4_* being the winning model of utility, suggest that subjects were sensitive to task cost timing, but much more idiosyncratic about preferences across the group. Collectively, subjects’ task preferences are idiosyncratic, but underlying these preferences is a consistent tendency to pre-crastinate physical effort.

### Choice Behavior Reflects Subjective Utility

One critique may be that the choice behavior is simply the result of a forced choice task, and subjects choose an arbitrary heuristic that is adhered to consistently. To further investigate this, we also characterized *how* subjects made and reported their choices. Specifically, we look at the deliberation time and the response vigor for each choice and how these measures correlate to the difference in subjective utility. If subjects chose arbitrarily, choice behavior may not correlate to the subjective utility used to model subject preferences.

Deliberation times varied across subjects (mean±s.e.: 0.629±0.127), yet consistently decrease as the difference in utility increases. For each model of subjective utility, we compare model fits for deliberation time and response vigor (Table 4). Across all models, we see a significant effect of difference in normalized subjective utility, where a negative slope indicates a decreasing (or quicker) deliberation time as the difference in subjective utility increases (*U_1_*: β **= −**0.152 [−0.172, −0.132], p<0.0001; *U_2_*: β **=** −0.181 [−0.204, −0.157], p<0.0001; *U_3_*: β = −0.178 [−0.200, −0.156], p<0.0001; *U_4_*: β **=** −0.206 [-0.231, −0.181], p<0.0001). Our data reflects the same trends as previously observed: easier choices are made more quickly (Festinger 1943).

**Table 4:**
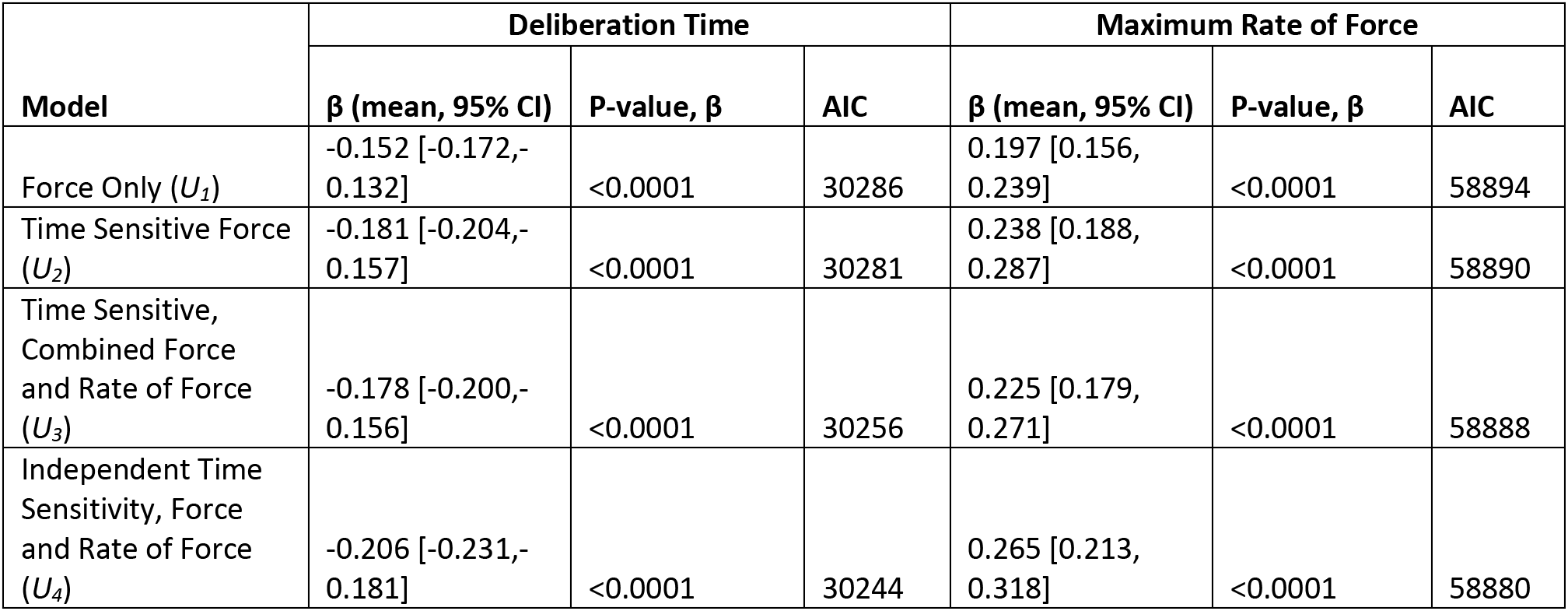
Model fit comparison for deliberation time and maximum cursor velocity across models of subjective utility.

| Model | Deliberation Time |  |  | Maximum Rate of Force |  |  |
| --- | --- | --- | --- | --- | --- | --- |
| | $\beta$ (mean, 95% CI) | P-value, $\beta$ | AIC | $\beta$ (mean, 95% CI) | P-value, $\beta$ | AIC |
| Force Only ($U_1$) | -0.152 [-0.172, -0.132] | <0.0001 | 30286 | 0.197 [0.156, 0.239] | <0.0001 | 58894 |
| Time Sensitive Force ($U_2$) | -0.181 [-0.204, -0.157] | <0.0001 | 30281 | 0.238 [0.188, 0.287] | <0.0001 | 58890 |
| Time Sensitive, Combined Force and Rate of Force ($U_3$) | -0.178 [-0.200, -0.156] | <0.0001 | 30256 | 0.225 [0.179, 0.271] | <0.0001 | 58888 |
| Independent Time Sensitivity, Force and Rate of Force ($U_4$) | -0.206 [-0.231, -0.181] | <0.0001 | 30244 | 0.265 [0.213, 0.318] | <0.0001 | 58880 |

Comparing models of deliberation time using AIC, we find that the fourth model of utility—the same model that best predicts choice behavior—best predicts deliberation time (AIC = 30244, minimum ΔAIC = 12, Figure 5A).

**Figure 5:**
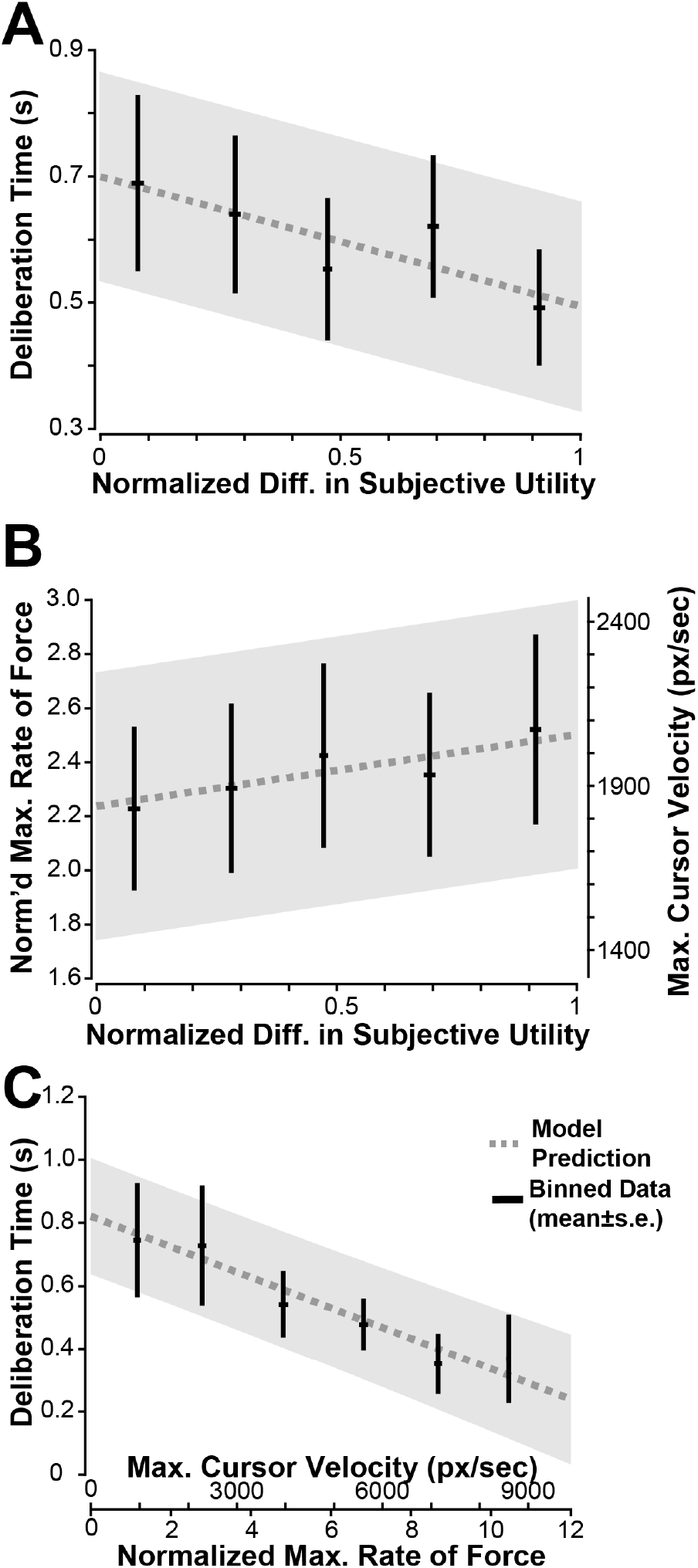
For the winning model of subjective utility (*U4*), **(A)** the mean and standard error for binned deliberation time (black) is plotted against the normalized absolute difference in subjective utility between force profiles. Deliberation time decreases as the difference in normalized subjective utility increases **(B)** The mean and standard error for binned choice vigor is plotted against the normalized difference in subjective utility between force profiles. Choice vigor is the maximum rate of force, manifested as the maximum cursor velocity for a choice. Choice vigor increases as the difference in normalized subjective utility increases. **(C)** As choice vigor increases, deliberation time decreases. Mean and standard error for deliberation time binned by choice vigor is shown in black. In all panels, the linear model prediction with 95% confidence intervals is shown in gray.

Interestingly, choices with a greater difference in utility were also made more vigorously. Choice vigor is fit using each model of utility with a linear mixed effects model that includes a random effect of subject. All models of choice vigor have significantly positive correlations to the difference in utility (*U_1_*: β **=** 0.197 [0.156, 0.239], p<0.0001; *U_2_*: β = 0.238 [0.188, 0.287], p<0.0001; *U_3_*: β**=** 0.225 [0.179, 0.271], p<0.0001; *U_4_*: β =0.265 [0.213, 0.318], p<0.0001). Again, we find that the fourth model best predicts choice vigor (AIC = 58880, minimum ΔAIC = 8, Figure 5B).

Independent of subjective utility models, we find that choices that were made faster (i.e., shorter deliberation time) were also made more vigorously (β **= −**0.0489 [−0.0588, −0.0390], p<0.0001), shown in Figure 5C.

In summary, variations in deliberation time and response vigor are best explained by the same model of subjective utility that best explains subject preferences, supporting the idea that choices were not just made with an arbitrary, consistently executed heuristic. Collectively, these results reinforce our findings that a subjective model of utility includes independently time-sensitive penalties on physical effort costs and task costs, and further suggest there is a link between an individual’s subjective valuation of utility, their decisions, and their movement control.

### Subjects Do Not Fatigue

We examined if subjects fatigued during the experiment. Using a linear model, we find that there are no significant effects of experiment phase (start and end of the *Familiarization Phase*: β=2.83×10^-3^,[−0.122, 0.127], p=0.964; start and end of the Session: β=0.119,[−0.00641, 0.244], p=0.0627, start and end of the *Choice Phase*: β=-0.116,[−0.00923, 0.241], p=0.0692), nor between hill or valley session types (β= −4.27×10^-3^,[−0.106, 0.0978], p=0.934) on subject’s maximum force exerted (Figure 6A). The lack of difference suggests that subjects are not decreasing in their ability to perform the task.

**Figure 6:**
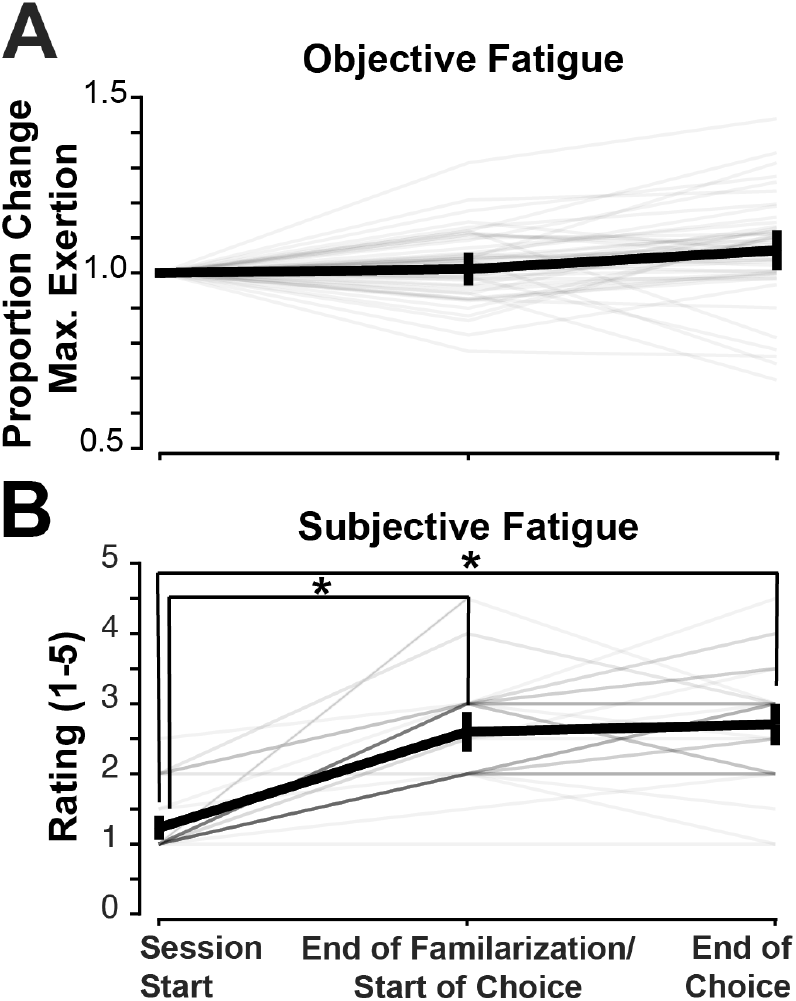
**(A)** Objective fatigue does not change throughout the experiment. The proportion of change of the maximum force exerted is plotted by phase. Subject means and standard error bars are shown in black with individual subjects plotted in gray. **(B)** Subjective fatigue ratings (1-5) are plotted by phase. Subject means and standard error bars are shown in black with individual subjects in gray. A significant effect of phase is indicated with asterisks, seen between the start of the session and the end of familiarization and start of the session and end of the full session.

Fatigue was also assessed via self-report using a Likert scale rating from one to five (Figure 6B). Subjects generally began with little to no fatigue (most rated their fatigue as a 1), with a significant increase in rating between the start and end of the *Familiarization Task* (β=1.37, [1.16, 1.58], p<0.0001) and start and end of the Session (β=1.48, [1.28, 1.69], p<0.0001), but no difference between start and end of the *Choice Task* (β=0.113, [−0.0945, 0.320], p =0.284) nor of session type (β=-0.158, [−0.327, 0.0108], p=0.0664). These results suggest that subjects did not fatigue objectively nor subjectively throughout the choice period, indicating that decision-making was not skewed or changing due to fatigue throughout a given session.

### Accuracy as a Predictor for Preference

All subjects performed the isometric arm-pushing tasks with less than the maximum 7% error. No subject failed a *Force Trial* in the *Familiarization Phase* nor the *Realization Trials* during the *Choice Phase*. During the *Familiarization Phase*, subjects performed with trials an error of 1.88%±0.180% (mean±s.e.), and in the *Choice Phase*, with an error of 2.35%±0.285% (mean±s.e.). In both phases, subjects exceeded the lenient accuracy requirements.

The difference in error was significantly related to the probability of choosing a profile (β=−44.5, [−49.4, − 39.6], p<0.0001), where the profile that is more accurately performed is preferred. However, the difference in duration had a significant effect on the difference in accuracy (β=1.68×10^-4^, [4.01×10^-5^, 2.96×10^-4^], p=0.0101), but the difference in onset time did not (β=-9.58×10^-6^,[−5.10×10^-5^, 3.18×10^-5^], p=0.650). Collectively, this suggests that the difference in hill or valley duration is likely the driver behind preference but happens to covary with accuracy.

## DISCUSSION

Here, we find that preferences for effortful isometric arm pushes are best explained by a combination of physical effort and task costs, both of which are time-sensitive. Using our approach, we separate physical effort and task costs and find that subjects significantly pre-crastinate physical effort, but are idiosyncratic about preference for when task costs occur. We propose a novel generalizable model of the time-sensitivity of effort that not only describes subject preferences but also choice deliberation time and vigor.

This study builds on a number of others investigating the subjective valuation of effort. While much work in biomechanics has demonstrated the importance of metabolic cost as a measure of physical effort, recent work has suggested the effort is more than metabolic cost alone. Effort appears to be subjectively valued with cost growing nonlinearly with energy or force (Hartmann et al. 2013; Morel et al. 2017; O’Brien and Ahmed 2019). Moreover, the subjective valuation varies idiosyncratically across individuals (Summerside and Ahmed 2021). Here we demonstrate that the time of effort investments also influence decision making over and above force or energy costs alone.

Our results are in family with many other pre-crastination studies, where a majority of subjects are pre-crastinators, but it is not unanimous (Blinch and DeWinne 2019; Fournier et al. 2019a; Kahneman et al. 1993; Patterson and Kahan 2020; Rosenbaum et al. 2014). We also see about two-thirds of our subjects pre-crastinate (16 of 25 subjects pre-crastinate physical effort). Currently, there are no conclusive measures that can stratify and predict who in a sampled population will decide to pre-crastinate or procrastinate. With further investigation, we may better understand the underlying mechanisms of pre-crastination beyond task-specific descriptors.

### Disentangling Physical Effort and Task Costs

The nature of physical effort makes it difficult to be independent from tasks costs. Naturally, most studies in pre-crastination of movement involve tasks that require physical effort to accomplish, like moving a bucket or transferring items (Fournier et al. 2019a, 2019b; Rosenbaum et al. 2014). Even cognitive tasks combine high mental workload before and during task completion (McBride et al. 2023; VonderHaar et al. 2019). Unique to our paradigm, we successfully tease apart high task costs and physical effort through the valley force profiles, where the task requires less physical effort.

Unsurprisingly, subjects were sensitive to the total amount of physical effort, in line with other studies that show that subjects prefer shorter durations of high force and lower total force overall (Körding et al. 2004). However, the time sensitivity results were surprising. If it were in line with current pre-crastination studies, we would expect to see pre-crastination of tasks costs, and perhaps time insensitivity of physical effort; however, we find the opposite. Why are subjects pre-crastinating physical effort?

### When task-independent, is physical effort just discomfort?

If physical effort is not required to complete a task, it may be valued similarly to pain. Physical effort, in this regard, is no longer an investment to receive a reward, but rather a standalone cost. In isometric arm-pushing tasks, subjects are shown to be sensitive to the duration of the physical effort (Hartmann et al. 2013; Körding et al. 2004). Holding a high level of force creates discomfort over time, so holding force for a shorter period of time near the finish ends on a “better note.” This explanation is in line with pain literature (Kahneman et al. 1993).

Unlike pain research, which relies on proprioception and is measured retrospectively, in this experiment, effort was accurately visually represented in a prospective decision-making task. If the choice task was redesigned so effort profiles were experienced sequentially without visual feedback, then a choice was made (similar to protocols found in Kahneman et al. 1993 and Körding et al. 2004), it may strengthen the observation of pre-crastination due to the peak-end rule.

### Is physical effort pre-crastinated due to a lack of uncertainty?

Within our paradigm, subjects accurately performed all profiles and did not fatigue, suggesting that subjects were confident in their ability to perform the required tasks. Additionally, force profiles were accurately visually represented, thus there was no uncertainty about what the task required. Could this certainty about the necessity to perform physical effort and the ability to do so, cause subjects to pre-crastinate?

Pre-crastination may be influenced by the certainty in which you must pay a cost. If temporal discounting of reward can be explained by the uncertainty about whether you will receive it in the future, what about the uncertainty of whether a cost must be paid in order to receive the reward? Consider a monetary example. A teenager has saved up enough money for a vehicle, but they are approaching their sixteenth birthday. Assuming they could not drive until they were of age, would they buy a car with their own money that they saved, or wait to possibly get that car as a gift? In other words, if there is uncertainty about a cost, without risking the chance of a reward, it should be procrastinated.

The same might be true with movement costs. If you knew the end of a walk was at the same elevation as its starting point, you wouldn’t voluntarily walk up a big hill. Rationally, you would try to stay as level as possible for as long as possible, procrastinating high physical effort. In the case of our study and many pre-crastination studies, the physical effort and task costs required are well-defined: a set of buckets is visible in front of the subject (Fournier et al. 2019a, 2019b; Rosenbaum et al. 2014), or in our case, a force profile is visually depicted to choose between.

In motor control, there is performance-based uncertainty. On a very short timescale where endpoint accuracy is required, one could argue that physical effort in saccades (Harris and Wolpert 1998) and reaches (Flash et al. 1992) are pre-crastinated. Velocity profiles are skewed due to signal-dependent noise and the speed-accuracy trade-off, where faster speeds are performed earlier and slower speeds, later to approach a target more accurately. When timing accuracy was required in a walking task, subjects also pre-crastinated (Tiew et al. 2020). It seems that if target accuracy is required in either space or time, a control strategy that pre-crastinates physical effort is favorable for task performance. Our paradigm does not involve target accuracy, and the effort invested throughout the profile is constrained. If this were the explanation for physical effort pre-crastination, it would suggest there is an underlying control strategy that extends across all forms of physical exertion.

Complementary to the uncertainty about needing to pay a cost, there is a tendency to procrastinate effort if there is uncertainty about the ability or willingness to perform it. In running racing, a constant pace is optimal (Behncke 1997; Keller 1973; Reardon 2013), and racers should expend all the energy they can. Yet a racer may be uncertain about how much capacity they have to expend energy near the end of the race. Countless racers have “bonked” due to a poor model of their remaining capacity. Did these racers over-precrastinate physical effort? Uncertainty about their capacity to perform the necessary work is typically manifested as runners racing more conservatively and procrastinating high effort until the end with a kick, sprinting to the finish when they are certain they have enough capacity remaining.

In our paradigm, physical effort costs are low, and subjects do not fatigue. Recent evidence suggests pre-crastination is the “automatic” response when costs are low, but as costs scale up, subjects care more about efficiency and begin to procrastinate. In Experiment 9 of Rosenbaum et al. 2014, when the weight of the buckets was increased, participants began to choose the bucket closer to the endpoint, minimizing physical effort and procrastinating task costs (Rosenbaum et al. 2014). In another experiment where subjects were asked to transport water cups of varying fullness and distances, when the cup was more full, requiring higher attentional resources not to spill (or task cost, in our words), subjects opted to procrastinate the task, carrying it a shorter distance. When the glass was less full, more subjects chose to pre-crastinate, picking up the cup closer to the subject and carrying it longer (Raghunath et al. 2021). As demand for either physical effort costs or task costs increases, these costs seem to become procrastinated. It would be interesting if we scaled up the physical effort in our paradigm, in either intensity or duration, whether we would see the same shift in preference towards procrastination.

Circling back to runners, consider a group that should have an accurate model of their energetic capacity: world record holders for the marathon. These runners pre-crastinate physical effort with a first half-marathon significantly faster than the second half (Díaz et al. 2018). Could pre-crastination of physical effort be about expending a limited resource like energy? Effort performed earlier with more capacity remaining could be subjectively preferable to effort performed later. This explanation offers an alternative hypothesis to some results, such as a bucket emptying task (Fournier et al. 2019b), where subjects chose to empty the heavier bucket first, and a dodgeball moving task (Rosenbaum and Dettling 2023), where subjects chose to carry larger loads of dodgeballs initially. Both found that subjects pre-crastinated expending physical effort, which a scarcity-based model of physical effort expenditure could explain. However, in these paradigms, physical effort is equated to subtask completion. The bigger task (like a heavier bucket) requires more physical effort, so this could still be about accomplishing the bigger subtask earlier. In contrast, our results suggest that physical effort is pre-crastinated, even when task effort is procrastinated. Because our subjects had high confidence in ability to perform the task, could this cause a pre-crastinating strategy?

Taken as a whole, the certainty of physical effort expenditure and the low magnitude of physical effort may be an explanation for why subjects tended to pre-crastinate. A possible direction for future studies is to manipulate uncertainty to see if there is a shift in time sensitivity.

### Choice Behaviors Reflect Utility

We show that choice behaviors reflect subjective utility. Similar to what is found in perceptual discrimination tasks, the more similar two stimuli are, the longer it takes to distinguish between them; thus, more difficult decisions are made more slowly (Festinger 1943). In our study, the closer in value the subjective utility is between two options, the longer it took subjects to make that choice. Another finding is that a change in the vigor of the response readout is also found with decision difficulty, in line with some recent studies (Smith and Peters 2022). Previous studies have shown saccades are made faster to more rewarding targets (Takikawa et al. 2002; Xu-Wilson et al. 2009), saccade vigor reflects the subjective utility of a decision when the effort is varied (Shadmehr et al. 2019). Only recently, Korbisch et al. have shown that a difference in utility between options is reflected in the vigor of saccades made during the decision-making process, where the greater the difference between the utility of choices, the greater the difference in peak saccade velocity between each option (Korbisch et al. 2022).

Here, we also show choice vigor varies with deliberation time. These results build on previous findings about vigor and temporal sensitivity, where individuals who made decisions about effortful tasks faster also made saccades more vigorously (Korbisch et al. 2022). Additionally, individuals who made saccades faster (Choi et al. 2014) and individuals who reached with greater vigor (Berret and Baud-Bovy 2022) had a higher cost on time. Using this metric, perhaps one can infer the time-sensitivity and preferences of an individual by how they make their choices.

### Limitations

One critique of our experiment design is that we sampled too coarsely. We intentionally test discrete values of hill and valley duration and onset time for test efficiency and the sake of between-subject comparison, but perhaps these were too coarsely sampled with a minimum of two-second differences in onset time and/or duration between profiles for a given choice trial. We do not find a well-defined indifference point between time-preference and duration, though we could predict where this might occur using our subjective utility model. Taking an adaptive sampling approach, we may have been able to define how much extra physical effort is worth performing at a given time more precisely.

A precise measure of objective cognitive effort remains unclear. In our formulation of utility, task costs may not map to cognitive effort. Using the rate of force squared as a representation of task costs is in line with a controls perspective, where a change of state requires control input. Force generation (i.e., a positive rate of force) is more metabolically costly than force maintenance (Russ et al. 2002), so, at least partially, our task costs could also be contributing to physical effort costs. Using working memory as an explanation, Rosenbaum et al. hypothesize that cognitive effort is high until the task is completed (Rosenbaum et al. 2014). This explanation also aligns with other research that holding a goal in mind is cognitively taxing (Einstein et al. 2003). Nevertheless, we could argue that the moment where greater attention is required, leading to higher cognitive effort, is during the act of picking up the bucket. This correlates well with our formulation that task costs are when changes in force occur.

## Conclusions

In this study, we designed an isometric arm-pushing task that can tease apart subjects’ preferences for physical effort, tasks costs, and time sensitivity. We find that subjects exhibit a preference for when both physical effort costs and task costs occur, adding to the body of evidence that costs in motor control are indeed time-sensitive. We find that, on average, subjects choose to pre-crastinate physical effort but are idiosyncratic about task costs. This study proposes a generalizable descriptive model of subjective utility, which predicts subject preferences as well as choice deliberation time and choice vigor.

## GLOSSARY

*α*: Weighting factor between cost terms, model parameter

*ψ*: Temporal sensitivity parameter

*τ*: Temperature parameter

*E*: Effort costs

*F*: Force, normalized to a subject’s maximum force exertion

*F*^•^: Rate of force,, normalized to a subject’s maximum force exertion

*K_physical_*: Normalizing constant for physical effort costs

*K_task_*: Normalizing constant for task costs

*p*: Predicted probability of preferring the left option

*R*: Reward

*s*: Temporal sensitivity function

*t*: Time elapsed from start to end of trial

*T*: Completion time of a force trial

*U*: Subjective utility

*y*: Binary outcome variable if the left option is chosen

## DATA AVAILABILITY

Source data for this study are openly available at https://github.com/neuromechanics-at-cu/precrastination_isometric_force.

## AUTHOR CONTRIBUTIONS

CMH and AAA conceived and designed research, edited and revised the manuscript, and approved the final version of the manuscript. CMH performed experiments, analyzed data, interpreted the results of experiments, prepared figures, and drafted the manuscript.

## REFERENCES

Alexander RMcN. A minimum energy cost hypothesis for human arm trajectories. Biol Cybern 76: 97– 105, 1997.

Bautista LM, Tinbergen J, Kacelnik A. To walk or to fly? How birds choose among foraging modes. Proc Natl Acad Sci 98: 1089–1094, 2001.

Behncke H. Optimization models for the force and energy in competitive running. J Math Biol 35: 375– 390, 1997.

Berret B, Baud-Bovy G. Evidence for a cost of time in the invigoration of isometric reaching movements. J Neurophysiol 127: 689–701, 2022.

Blinch J, DeWinne CR. Pre-crastination and procrastination effects occur in a reach-to-grasp task. Exp Brain Res 237: 1129–1139, 2019.

Choi JES, Vaswani PA, Shadmehr R. Vigor of Movements and the Cost of Time in Decision Making. J Neurosci 34: 1212–1223, 2014.

Cohen RG, Rosenbaum DA. Where grasps are made reveals how grasps are planned: generation and recall of motor plans. Exp Brain Res 157: 486–495, 2004.

Díaz JJ, Fernández-Ozcorta EJ, Santos-Concejero J. The influence of pacing strategy on marathon world records. Eur J Sport Sci 18: 781–786, 2018.

Einstein GO, McDaniel MA, Williford CL, Pagan JL, Dismukes RK. Forgetting of intentions in demanding situations is rapid. J Exp Psychol Appl 9: 147–162, 2003.

Festinger L. Studies in decision: I. Decision-time, relative frequency of judgment and subjective confidence as related to physical stimulus difference. J Exp Psychol 32: 291–306, 1943.

Flash T, Inzelberg R, Schechtman E, Korczyn AD. Kinematic analysis of upper limb trajectories in Parkinson’s disease. Exp Neurol 118: 215–226, 1992.

Fournier LR, Coder E, Kogan C, Raghunath N, Taddese E, Rosenbaum DA. Which task will we choose first? Precrastination and cognitive load in task ordering. Atten Percept Psychophys 81: 489–503, 2019a.

Fournier LR, Stubblefield AM, Dyre BP, Rosenbaum DA. Starting or finishing sooner? Sequencing preferences in object transfer tasks. Psychol Res 83: 1674–1684, 2019b.

Goldman MD, Marrie RA, Cohen JA. Evaluation of the six-minute walk in multiple sclerosis subjects and healthy controls. Mult Scler J 14: 383–390, 2008.

Haith AM, Reppert TR, Shadmehr R. Evidence for Hyperbolic Temporal Discounting of Reward in Control of Movements. J Neurosci 32: 11727–11736, 2012.

Harris CM, Wolpert DM. Signal-dependent noise determines motor planning. Nature 394: 780–784, 1998.

Hartmann MN, Hager OM, Tobler PN, Kaiser S. Parabolic discounting of monetary rewards by physical effort. Behav Processes 100: 192–196, 2013.

Hull CL. Principles of behavior: an introduction to behavior theory. Oxford, England: Appleton-Century, 1943.

Kahneman D, Fredrickson BL, Schreiber CA, Redelmeier DA. When More Pain Is Preferred to Less: Adding a Better End. Psychol Sci 4: 401–405, 1993.

Keller JB. A theory of competitive running. Phys Today 26: 43–47, 1973.

Klein-Flügge MC, Kennerley SW, Saraiva AC, Penny WD, Bestmann S. Behavioral Modeling of Human Choices Reveals Dissociable Effects of Physical Effort and Temporal Delay on Reward Devaluation. PLOS Comput Biol 11: e1004116, 2015.

Korbisch CC, Apuan DR, Shadmehr R, Ahmed AA. Saccade vigor reflects the rise of decision variables during deliberation. Curr Biol 32: 5374–5381.e4, 2022.

Körding KP, Fukunaga I, Howard IS, Ingram JN, Wolpert DM. A Neuroeconomics Approach to Inferring Utility Functions in Sensorimotor Control. PLOS Biol 2: e330, 2004.

Lamb DG, Correa LN, Seider TR, Mosquera DM, Rodriguez JA, Salazar L, Schwartz ZJ, Cohen RA, Falchook AD, Heilman KM. The aging brain: Movement speed and spatial control. Brain Cogn 109: 105– 111, 2016.

Mazzoni P, Hristova A, Krakauer JW. Why Don’t We Move Faster? Parkinson’s Disease, Movement Vigor, and Implicit Motivation. J Neurosci 27: 7105–7116, 2007.

McBride DM, Villarreal SR, Salrin RL. Precrastination in cognitive tasks. Curr Psychol 42: 14984–15002, 2023.

Morel P, Ulbrich P, Gail A. What makes a reach movement effortful? Physical effort discounting supports common minimization principles in decision making and motor control. PLOS Biol 15: e2001323, 2017.

Myerson J, Green L. Discounting of delayed rewards: Models of individual choice. J Exp Anal Behav 64: 263–276, 1995.

O’Brien MK, Ahmed AA. Asymmetric valuation of gains and losses in effort-based decision making. PLOS ONE 14: e0223268, 2019.

Patterson EE, Kahan TA. Precrastination and the cognitive-load-reduction (CLEAR) hypothesis. Memory 28: 107–111, 2020.

Raghunath N, Fournier LR, Kogan C. Precrastination and individual differences in working memory capacity. Psychol Res 85: 1970–1985, 2021.

Ralston HJ. Energy-speed relation and optimal speed during level walking. Int Z Für Angew Physiol Einschließlich Arbeitsphysiologie 17: 277–283, 1958.

Reardon J. Optimal pacing for running 400- and 800-m track races. Am J Phys 81: 428–435, 2013.

Redelmeier DA, Kahneman D. Patients’ memories of painful medical treatments: real-time and retrospective evaluations of two minimally invasive procedures. Pain 66: 3–8, 1996.

Rigoux L, Guigon E. A Model of Reward- and Effort-Based Optimal Decision Making and Motor Control. PLOS Comput Biol 8: e1002716, 2012.

Rigoux L, Stephan KE, Friston KJ, Daunizeau J. Bayesian model selection for group studies — Revisited. NeuroImage 84: 971–985, 2014.

Rosenbaum DA, Dettling J. Carrying groceries: More items in early trips than in later trips or the reverse? Implications for pre-crastination. Psychol Res 87: 474–483, 2023.

Rosenbaum DA, Gong L, Potts CA. Pre-Crastination: Hastening Subgoal Completion at the Expense of Extra Physical Effort. Psychol Sci 25: 1487–1496, 2014.

Rosenbaum DA, Sauerberger KS. Deciding what to do: Observations from a psycho-motor laboratory, including the discovery of pre-crastination. Behav Processes 199: 104658, 2022.

Russ DW, Elliott MA, Vandenborne K, Walter GA, Binder-Macleod SA. Metabolic costs of isometric force generation and maintenance of human skeletal muscle. Am J Physiol-Endocrinol Metab 282: E448–E457, 2002.

Schwob N, Epping A, Taglialatela J, Weiss D. Animal Behavior and Cognition The Early Bonobo Gets the Juice? The Evolutionary Roots of Pre-crastination in Bonobos (Pan paniscus). Anim Behav Cogn 9: 3–13, 2022.

Shadmehr R, Huang HJ, Ahmed AA. A Representation of Effort in Decision-Making and Motor Control. Curr Biol 26: 1929–1934, 2016.

Shadmehr R, Orban de Xivry JJ, Xu-Wilson M, Shih T-Y. Temporal Discounting of Reward and the Cost of Time in Motor Control. J Neurosci 30: 10507–10516, 2010.

Shadmehr R, Reppert TR, Summerside EM, Yoon T, Ahmed AA. Movement Vigor as a Reflection of Subjective Economic Utility. Trends Neurosci 42: 323–336, 2019.

Smith E, Peters J. Motor response vigour and visual fixation patterns reflect subjective valuation during intertemporal choice. PLOS Comput Biol 18: e1010096, 2022.

Stephan KE, Penny WD, Daunizeau J, Moran RJ, Friston KJ. Bayesian model selection for group studies. NeuroImage 46: 1004–1017, 2009.

Summerside EM, Ahmed AA. Using metabolic energy to quantify the subjective value of physical effort. J R Soc Interface 18: 20210387, 2021.

Takikawa Y, Kawagoe R, Itoh H, Nakahara H, Hikosaka O. Modulation of saccadic eye movements by predicted reward outcome. Exp Brain Res 142: 284–291, 2002.

Tiew EH, Seethapathi N, Srinivasan M. Pre-crastination: time uncertainty increases walking effort. bioRxiv: 2020.07.17.208140, 2020.

VonderHaar RL, McBride DM, Rosenbaum DA. Task order choices in cognitive and perceptual-motor tasks: The cognitive-load-reduction (CLEAR) hypothesis. Atten Percept Psychophys 81: 2517–2525, 2019.

Wasserman EA, Brzykcy SJ. Pre-crastination in the pigeon. Psychon Bull Rev 22: 1130–1134, 2015.

Xu-Wilson M, Zee DS, Shadmehr R. The intrinsic value of visual information affects saccade velocities. Exp Brain Res 196: 475–481, 2009.

